# Modelling post-disturbance empirical patterns in a forest ecosystem

**DOI:** 10.1101/2024.08.01.606184

**Authors:** Davide Zanchetta, Amos Maritan, Sandro Azaele

## Abstract

We investigate the impact of disturbances on forest ecosystems by examining transient population dynamics in a controlled experiment carried out in a tropical rainforest. We first model the mean species abundance with a simple consumer-resource model, which is then extended into two multi-species frameworks for community dynamics: a neutral model, which emphasises the ability of any species to recover independently of the others; and a non-neutral one, where species interactions play an important role in reconstructing ecosystem’s structure and patterns. The results indicate that, while both frameworks accurately describe correlation functions and mean-variance relations, the non-neutral model more effectively captures community structure as revealed by an evenness indicator. This suggests that interspecific interactions significantly influence the ecosystem’s ability to respond to disturbances, providing deeper insights into the recovery dynamics of forest ecosystems.

## INTRODUCTION

A substantial portion of ecological theory and our comprehension of ecological systems have traditionally been grounded in the notion that the states and dynamics observed in these systems align with their stable asymptotic behavior. This traditional focus on the long-term predictions of mathematical models contrasts with the more immediate time frames often considered in ecological observations and management practices [1–3].

Understanding how populations, communities, and ecosystems react to rapid, human-induced environmental changes is a significant challenge due to the diverse nature of disturbances, whose characteristics vary widely in aspects such as duration, spatial extent, intensity, frequency, and type. For instance, when considering duration, we may face two extreme cases such as pulse perturbations, describing disruptive but short-lived events, or press perturbations, which are milder but last for a longer time. As for spatial scales, perturbations may range from local scales (when due to events which are localized in space) to regional or even global ones (e.g. climate change) [4–6]. Ecological literature has shown that stability is a multifaceted and intrinsically multidimensional concept, with various components responding differently to different types of perturbations [7–11], resulting in a complex landscape of transient and asymptotic behaviors.

Investigating how an ecosystem evolves over time following a sudden disturbance necessitates moving beyond dynamics around steady states, which are more straightforward to analyze. However, even with minimal models, for which one derives analytic predictions for macroecological patterns (e.g. species abundance distribution), it is difficult to study states far from stationarity, given that even quite simple models do not have an explicit solution. Some steps in this direction have been taken for mathematically simple models [12, 13]. Moreover, a full comparison between data and models in the context of transient dynamics is also difficult. This gap exists both due to analytical difficulties and because data are often collected when the ecosystem of interest is supposedly at stationarity. Disturbances, if present, are not controlled and therefore are indistinguishable from fluctuations caused by endogenous processes. All these factors present substantial obstacles to our understanding of how perturbations impact large ecosystems.

In the following, on the contrary, we focus on a controlled forest disturbance experiment and develop two contrasting modelling frameworks for investigating how perturbations reverberate on the populations and structure of the ecosystem. We will look into a census data [14–16] in which several plots of a neotropical rainforest are subject to culling treatments of different intensity. Indicators of ecological dynamics (e.g. mean abundances and evenness) show non-monotonic trajectories or long relaxation times in the aftermath of disturbances (see Fig. 1). While estimates of above-ground biomass show monotonic return to a pre-disturbance level (see Fig. 3 of [17]), the dynamics of populations’ sizes might more directly affect long term biodiversity, as a result of individual-level processes and species interactions.

**FIG. 1:**
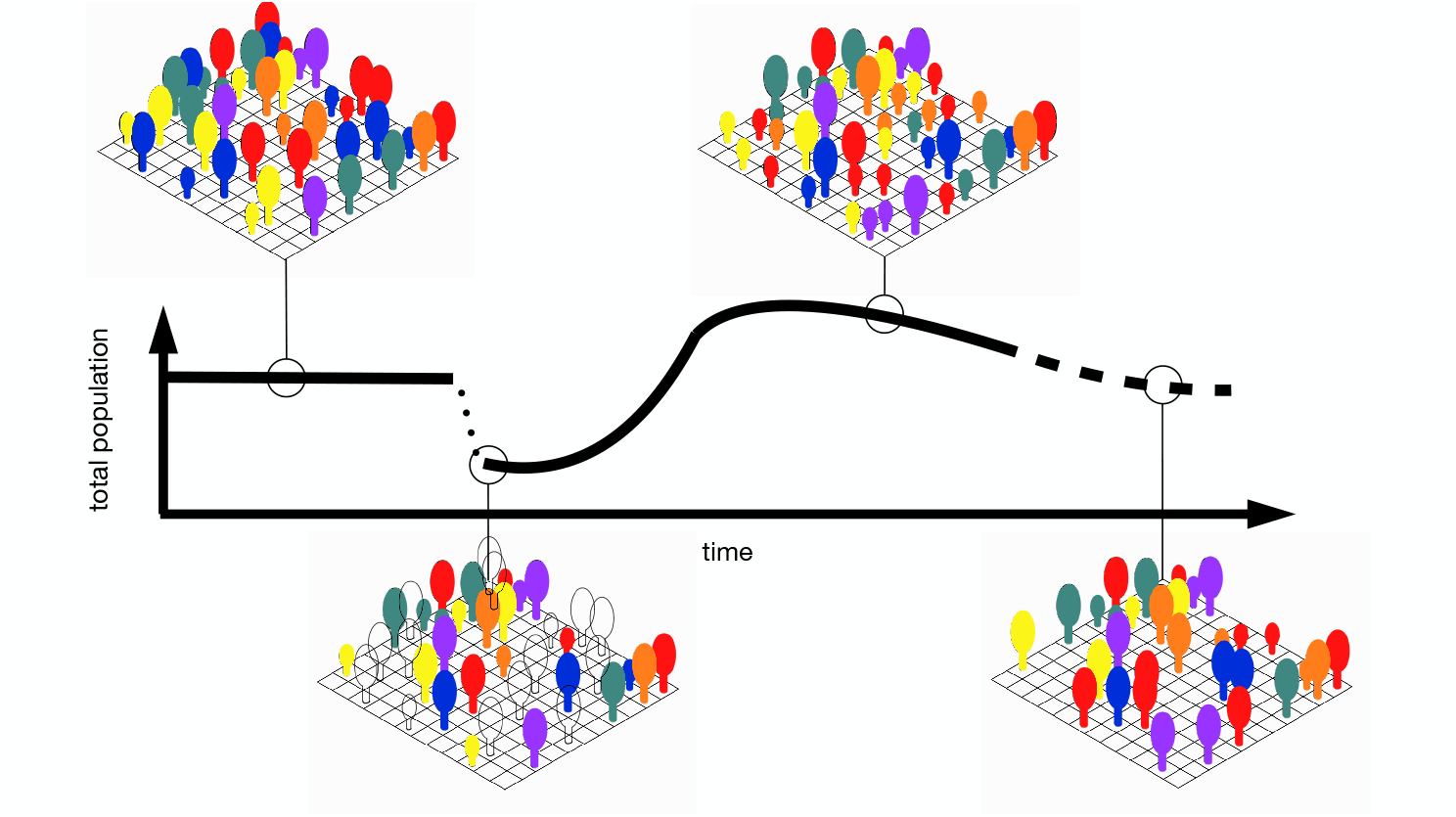
Population dynamics in a perturbed woodland. A mature forest stand at stationarity is decimated, with the most abundant species being the most affected; resources are freed up, enhancing recruitment at successive stages and leading to a total population that overshoots the initial average population; eventually, competition for resources drives the total population towards a sustainable value.

Neutral and non-neutral models have the ability to offer complementary insights into how disturbances impact ecological systems. The first ones [18, 19], which assume species equivalence and emphasize the importance of demographic random events in shaping community structure, can capture aspects which affect all species uniformly; providing a baseline for understanding ecosystem recovery driven by processes that transcend species’ identities and couplings. Deviations from neutral models’ predictions highlight the role of species-specific traits and interactions in response to disturbances. In contrast, non-neutral frameworks emphasize the role of competition, predation, and mutualism, in shaping community structure and dynamics. Here, habitat variability, different types of interactions or trade-offs in utilising resources may have an impact on differential responses to disturbances [20–23]. In this paper we leverage on these two complementary approaches to offer a more comprehensive understanding of ecosystem resilience and recovery.

Our aim is to model a forest stand that, starting from stable conditions, undergoes a substantial decrease in population size, which, however, does not lead to the eventual collapse of the whole community. Observations suggest that the removal of individuals creates additional space and resources, facilitating greater recruitment in subsequent stages. As a result, the population may eventually overshoot the initial stable size as indicated in Fig. 1. This behavior has been associated with ecosystem reactivity [24] and is important in transient dynamics. We will model this type of reaction to perturbation going beyond the linear regime. Notice that this behavior cannot be explained by naїve population models (namely, the population size cannot be governed by a first order local ODE), including the logistic model, which entails a monotonic time evolution after an initial value. More aptly, it suggests that a consumer-resource dynamics may be a more suitable modelling framework. Thus we start with a simple consumer-resource model which is able to capture the evolution of the average population size in the aftermath of a pulse perturbation, after which the mean abundance per species decreases by more than 30%, as in the experiment. We then refine this baseline model in two directions along the lines that we have alluded to above: on the one hand, the neutral dynamics of the community is described by a stochastic consumer-resource model in which the population and the resource are affected by multiplicative noises; this allows to keep track of the evolution of the emergent patterns of the community. On the other hand, we extend the baseline model with non-neutral elements by incorporating pairwise interactions between species; species-specific couplings are heterogeneous and structureless (quenched disorder) as assumed in more recent approaches [25–29]. The reason for this is to maintain a minimal level of complexity in the non-neutral model, while also ensuring a fair comparison with the neutral one.

We investigate the phenomenology of these two consumer-resource models following an initial sudden disturbance. We focus on the dynamics of community structure, including the evolution of species abundances, temporal correlation functions, evenness, and Taylor’s law. We compare these models to empirical data using various statistical measures within a Bayesian framework, allowing for a comprehensive evaluation of the two modelling frameworks.

## MATERIALS AND METHODS

### Dynamics of mean abundances

We first define the baseline model for the dynamics of the average abundance of species in the forest community. We use a consumer-resource model with two coupled variables, one describes the mean population size and the other one an average abiotic resource which supports species’ population. Whilst we have access to empirical data on the population dynamics, information on resources is lacking, hence we will consider it as a hidden degree of freedom. Besides being relatively simple, this approach offers the benefit of presenting a broad spectrum of behaviors in temporal trajectories, including overshooting curves in transient dynamics as displayed in empirical data. Let *N*_*i*_(*t*) be the number of individuals of species *i* (with *i* ∈ {1, …, *S*}) at time *t*, we indicate the mean abundance with 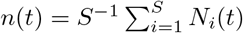and the mean resource with *c*(*t*), so the model reads

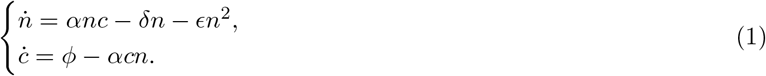

Consumers’ population increases with a per-capita rate *αc* and have a per-capita death rate *δ*, while a unit of resource is consumed at a rate *αn* and restored at a rate *ϕ/c*. We have also introduced a quadratic term in the equation for consumers with an effective self-interaction coupling *ϵ*. This is appropriate when considering spatial dynamics implicitly or as a consequence of interaction with pathogens [27]. We denote the initial conditions for consumers and resources as (*n*(0), *c*(0)) ≡ (*n*_0_, *c*_0_). For negative times the system is supposed to be at stationarity with (*n*(*t*), 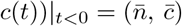, where

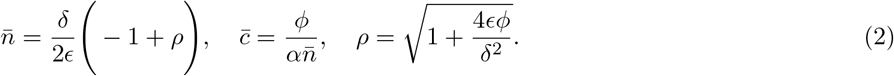

is the non-trivial stationary point of equations (1). At time *t* = 0 a pulse perturbation causes an abrupt decrease in the total population, quantified by 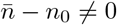. This difference represents the magnitude of the disturbance, which is generally significant. Notice that we suppose that the parameters of the model are not changed by the disturbance, which affects only the initial conditions. This implicitly assumes that the characteristic demographic scales of species are much larger than those of the human-induced change.

Further, linear stability analysis shows that a small perturbation near the equilibrium point eventually settles back into it with dampened oscillations when either 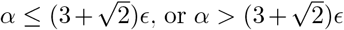 and *ϕ <* 4*αδ*^2^(*δ* ‐ *ϵ*)^2^*/*(*α*^2^ ‐ 6*αϵ* + *ϵ*^2^)^2^. The oscillatory behavior, as shown in Fig. 2 and discussed later, is essential for accurately representing the temporal trend of the data. The linearised dynamics around the fixed point gives an approximate solution which, however, does not match the full solution with sufficient accuracy to predict community patterns as we suggest in the following. Similar conclusions can be drawn when assuming a separation of time scales between *n* and *c*. In this scenario, one can deduce that the solution reaches stationarity monotonically over time (See Appendix B). However, in Appendix B we derive a uniform approximation to the full solution which better describes the temporal dynamics. This is expressed as

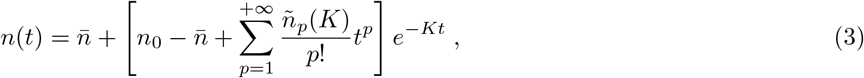

where 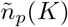 is defined in Eq.(2) and 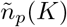 are coefficients that depend only on the initial conditions and the parameters of the model in Eq.(1), namely,

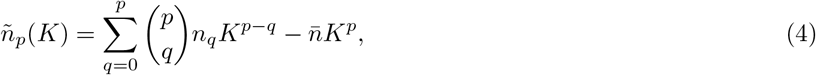

where 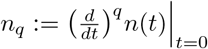, which depend only on the parameters of the system and the initial conditions. Because for *t*≫*K*^*‐*1^ the solution settles to its equilibrium value, 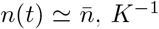 provides a characteristic temporal scale which measures how quickly the systems responds to and assimilates the perturbation. *K* can be roughly estimated by the smallest temporal rate of (1); however, when the parameters of the models are fitted to the data, the three independent temporal rates – *δ, δα/ϵ* and *ϕϵ/δ* – turn out to be of the same order of magnitude (see Table II). Therefore, *K* can be better considered a free parameter to be determined via non-linear regression with empirical data. A similar expansion can be also derived for *c*(*t*). In the next section, we extend (1) to predict trajectories of different species.

**FIG. 2:**
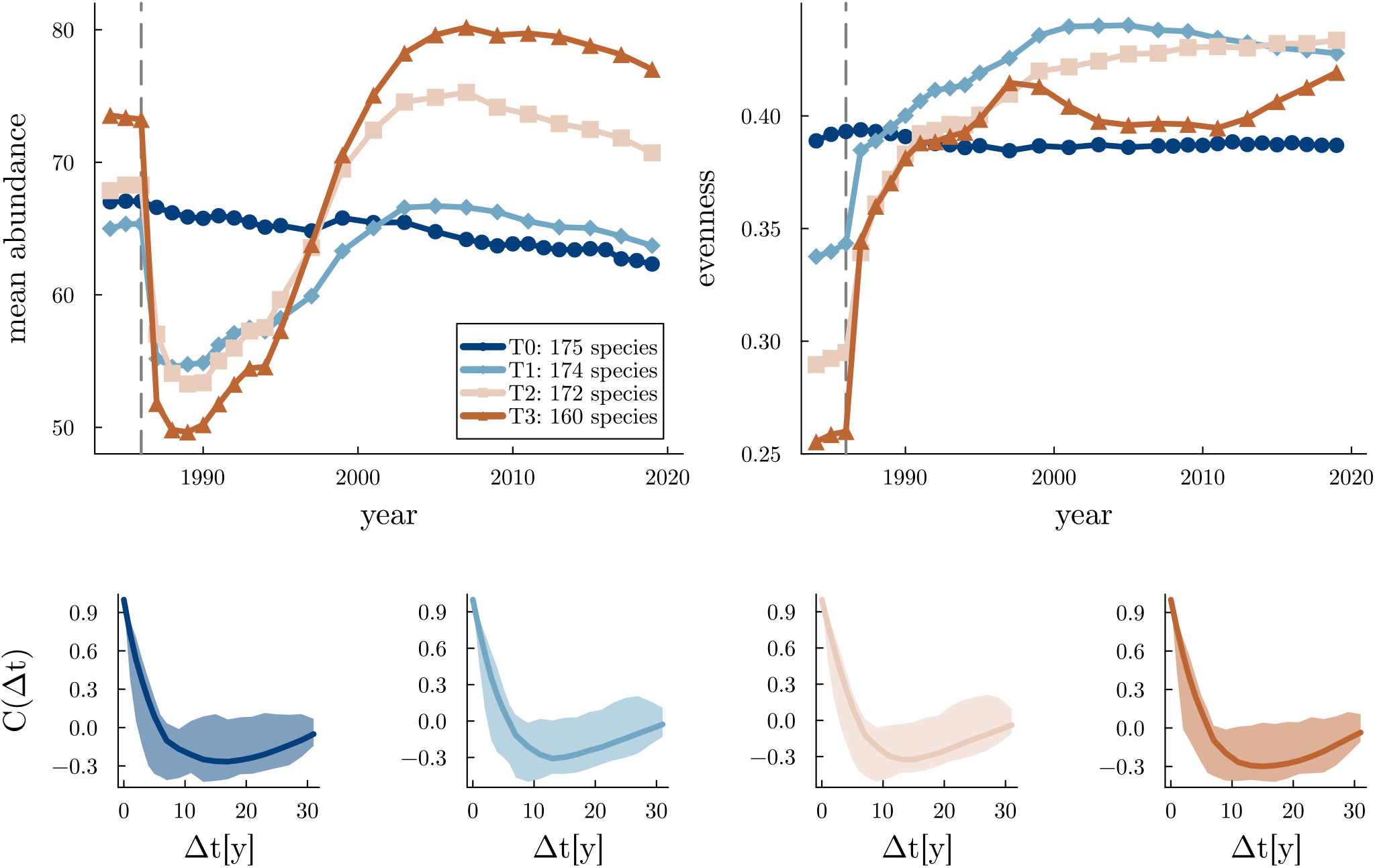
Representation of data from PDEx through quantities computed on different plot types. Top left: time series for mean abundances. Top right: time series for evenness. Bottom: temporal correlation function, with envelopes indicating 5%-95% quantiles, with correlation times reported on each plot. Different plot types are identified by different colors according to the legend of the top left image, i.e. dark blue for T0, light blue for T1, beige for T2 and brown for T3.

**FIG. 3:**
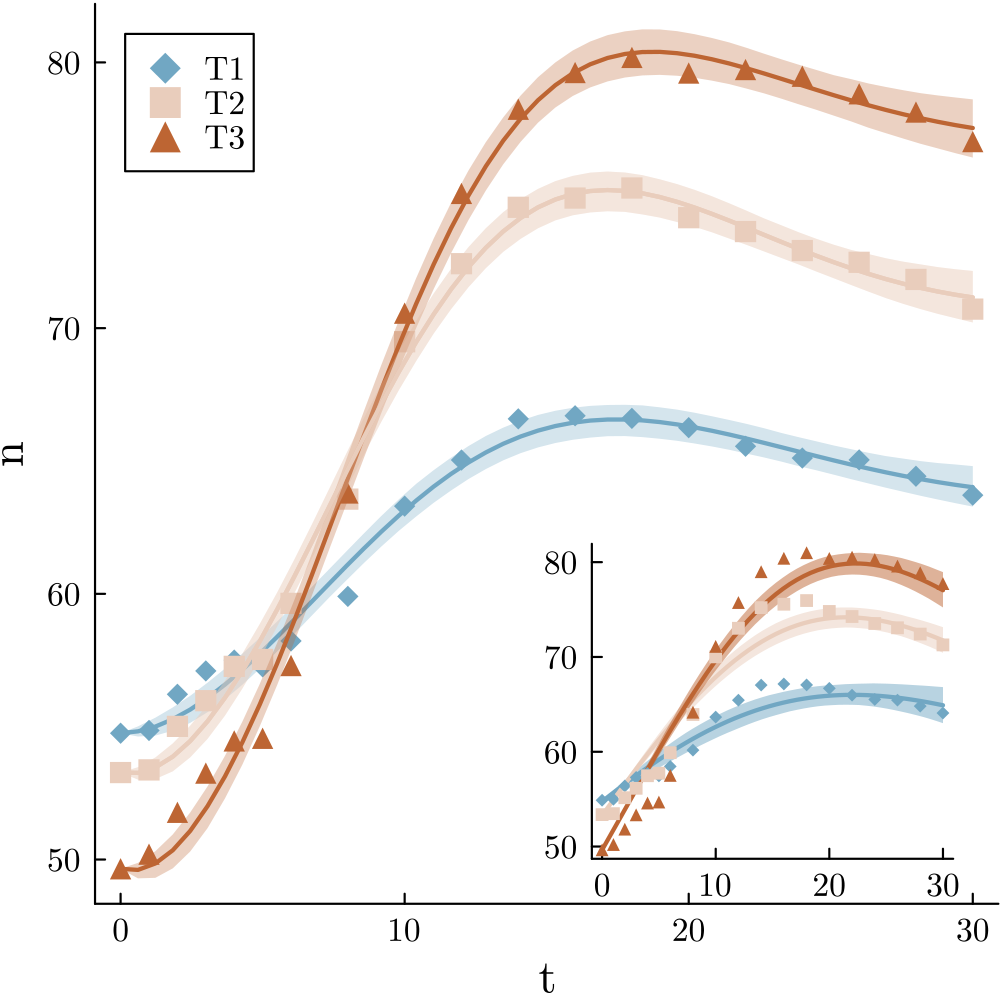
Mean abundances. Filled points represent empirical data and solid lines are means calculated from model runs (envelopes are 5-95% quantiles) with parameters which are sampled from the posterior distributions. Main panel represents fit of model (1) to each type of perturbed plots individually (i.e. the output is three sets of parameters); inset represents inference on all perturbed plot (i.e. the output is one set of parameters). Parameters are found in SI, table II.

### Multi-species dynamics

The model for mean abundances is further developed in the following sections to accommodate community dynamics, offering alternative strategies for restoring the initial state. The main point here is to explore whether recovery from disturbance is a process that involves the entire ecosystem, and therefore species’ interactions are important, or whether species can mainly recover on their own, i.e., independently of how other species evolve. In the latter case, we expect that a neutral approach should be able to account for the post-disturbance evolution, where random dispersal or demographic stochasticity play a crucial role. Neutral models assume functional equivalence at the individual level [30], which means that organisms in an ecosystem are demographically identical in terms of effective vital rates. This approach emphasizes the role of stochasticity in the recovery process, whilst species’ identity is a relatively minor factor.

If post-disturbance recovery involves the entire ecosystem, it is more appropriate to include interaction effects in the modeling framework to acknowledge the ecological non-equivalence of species. This approach underscores that species differ in their responses to environmental conditions, resource utilization, and interactions with other species. For instance, the way different species react to pathogens can lead to diverse interactions between them, resulting in a range of responses to external disturbances [27]. We then consider an alternative non-neutral model with heterogeneous and structureless interactions, namely, where these latter are drawn from a specific distribution. Ecological models with this type of interactions (which are known as quenched noise or disordered interactions in the physics literature), and their reduction to neutral models, have been extensively studied by statistical physicists [25, 26, 28, 29, 31], but to our knowledge, they have not been directly compared to transient dynamics observed in empirical data.

#### Neutral model

The neutral model is defined in a phenomenological manner by incorporating noise terms into the deterministic equations (1) to represent stochastic effects. In this framework, species’ population sizes are different realizations of a stochastic process. This allows us to calculate the species abundance distribution after a disturbance, as well as other important patterns such as the time correlation function and evenness. We use multiplicative noise terms, meaning that the noise strength depends on the abundances of consumers and resources. Because the noise term for consumers arises from individual birth and death events, it takes the form of demographic noise (namely, proportional to 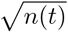 [32–35]). Instead, the noise term for abiotic resources originates from random variations in environmental parameters and therefore takes the form of environmental noise (that is, proportional to *c*(*t*) [32, 36]). Both these forms of multiplicative noise ensure that the variables *n*(*t*) and *c*(*t*) remain always non-negative. The model is formulated as two coupled stochastic differential equations.

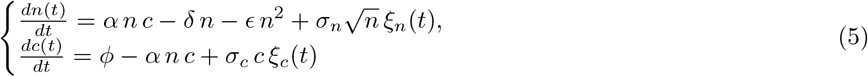

where *σ*_*n*_ and *σ*_*c*_ are the strengths of the demographic and environmental noises, respectively. Initial conditions for *n* are sampled from the distribution of abundances immediately after the perturbation, while initial condition for *c* is a fixed constant. We chose reflecting boundary conditions at *n* = 0 and *c* = 0, which effectively prevent species from disappearing entirely from the ecosystem, consistently with the fact that experimental plots are surrounded by (unperturbed) forest stands. More details can be found in Appendix C.

#### Non-neutral model

In models that are not neutral, species may exhibit various resource uptake abilities and respond differently to disturbances due to specific traits and interactions with neighboring species. Our modeling approach here does not focus on any particular trait; rather, we introduce interactions in a minimal way, that is, by assuming that they are heterogeneous, with no specific structure (quenched disordered interactions), and reduce to (1) as a model of mean abundances. It can be demonstrated that heterogeneous interactions are capable of mediating spatial effects, presence of pathogens, and other environmental factors [27, 37, 38] in a simple way. We break neutrality by replacing *ϵ* with a matrix of pair-dependent interactions, Λ_*ij*_, whose entries are random variables distributed as specified below. Because any species *i* has a population *N*_*i*_(*t*) and we define 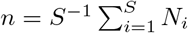, from (1) we find an *S* + 1-dimensional system of ordinary differential equations,

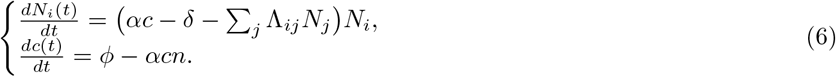

This model can be obtained from a consumer-resource dynamics with *S* species and *R* resources by averaging over resources (see Appendix A); we used this approach to reduce the number of free parameters. The first equation describes how the population size of the species *i* changes over time due to its interaction with species *j* and with an average resource density *c*. If we require that the average population size, 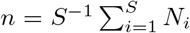, is exactly governed by equations (1), we obtain the conditions Λ_*ii*_ = *ϵ/S* and Λ_*ij*_ + Λ_*ji*_ = 2*ϵ/S*. This implies that species *i* and *j* interact in a very specific way, with fine-tuned correlations, which may be unrealistic. Thus, we relaxed those conditions and assumed that they are satisfied on average only; this can be obtained by defining that the entries of the interaction matrix Λ are

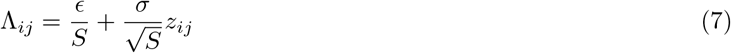

where *z*_*ij*_ are normal and uncorrelated random variables with mean 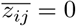 and variance 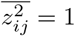. In the following we will investigate the effect of correlations between *z*_*ij*_ and *z*_*ji*_, i.e.,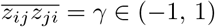. While the scaling 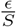 is necessary for recovering (1), the dependence on 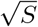 in (7) – which is customary in the physics literature of disordered systems – is not important in the current model, because we never take the limit *S*→ +∞ and we never compare communities with a different number of species. In our case *S* is a fixed number which corresponds to the empirical number of species of the ecosystem. In our phenomenological approach, all species possess equal capabilities in extracting nutrients from a single effective resource. Of course, this is an approximation, which helps, however, to manage the number of parameters for a fair comparison with the neutral model, which has the same number of free parameters. Also, it facilitates the comparison with the empirical data, because we use the parameters we obtained from the fit to (1), which only entails average population sizes. The heterogeneity of the interactions is controlled by *σ*, which we fix by looking at the time series of all species before the disturbance (see Appendix D for details). We find that *σ*≪ *ϵ*, so that most species interact weakly and almost always negatively affect each other because *ϵ >* 0. Therefore, interactions are weak, asymmetric and have mixed signs as was found in other forest stands [39].

In the next section, we look at the empirical data on forest ecosystem dynamics after a controlled disturbance, which provides a test for the models we have described.

## Data

Here we describe the empirical data from the Paracou Disturbance Experiment (PDEx) [15, 40]. The PDEx has been carried out in French Guiana since 1984. Twelve square plots of forest have been selected and destined for different treatments: 3 control plots were left undisturbed, while the others 9 were perturbed with different intensities, resulting in progressively more severe culling of populations across different species, with up to 56% of above-ground biomass lost. The culling treatments, which took place in 1986, are labelled according to their intensity as reported in the Tab. I; see Fig. 2, top row. In the following we use these labels to refer to different groups of plots, e.g. all plots which perturbed with treatment T3 will be referred to as ‘T3 plots’. The decrease in population occurred over roughly two years, so we model these disturbances as pulse perturbations.

**TABLE I:**
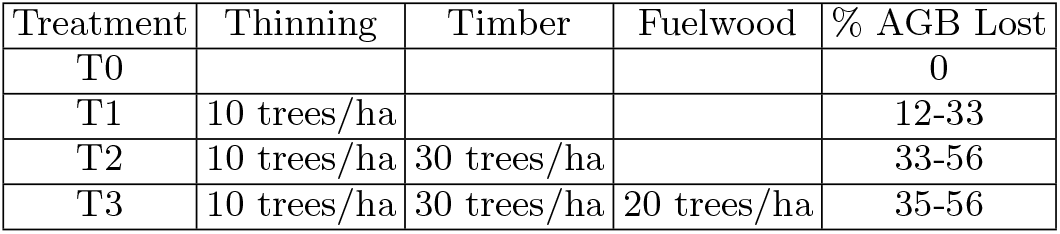
Table of treatments. Timber was collected from trees of DBH ≥ 50 or 60 cm, according to species; thinning was applied to all non-valuable trees of DBH ≥ 40 cm; fuelwood was collected from all trees for which 40 cm ≤ DBH ≤ 50 cm. For details, see [15].

| Treatment | Thinning | Timber | Fuelwood | % AGB Lost |
| --- | --- | --- | --- | --- |
| T0 |  |  |  | 0 |
| T1 | 10 trees/ha |  |  | 12-33 |
| T2 | 10 trees/ha | 30 trees/ha |  | 33-56 |
| T3 | 10 trees/ha | 30 trees/ha | 20 trees/ha | 35-56 |

Data collected contain, among other fields, the identity of each tree and its position within its plot, its diameter at breast height (DBH) and taxonomic identification, as well as its status (alive, subjected to treatment, dead, and so on); being interested in population dynamics without explicit representation of spatial effects, we only use trees identification code to track individual life events (birth and death), and species identification. The number of species exceeds 150 in all plot types. We focus on temporal dynamics of populations’ sizes, without considering invasion dynamics. Only trees with DBH *>* 10cm are censused, so young individuals are removed.

We pooled the populations of plots which received the same treatment and computed the mean abundance of species at each census year (see Fig. 2, top left). Following the disturbance, the number of individuals increases, exceeding the initial level after about 15 years.

To assess the effect of treatment on the temporal dynamics of community structure, we quantified for each year the evenness of each plot type community as follows. For an ecological community of *S* species represented by (*N*_1_, …, *N*_*S*_) individuals, the relative abundances of species are defined as *p*_*i*_ = *N*_*i*_*/*Σ _*j*_ *N*_*j*_; these are the probabilities that an individual drawn uniformly at random from the ecosystem belongs to species *i*. We define the evenness of a community with relative abundances *p* = {*p*_*i*_} _*i*_ as exp(*H*(*p*))*/S*, where *H*(*p*) =− Σ _*i*_ *p*_*i*_ log *p*_*i*_ is the Shannon entropy of the probability distribution *p* [41]. Our definition of evenness, which is one of many possible choices [42], is convenient because it separates species richness from community structure [43], thus allowing for the comparison of different communities. According to this definition, evenness assumes values between 1/S – when a species is much more abundant than the others – and 1 – which corresponds to all species having comparable abundances. All treatments affected more heavily the most abundant species, which resulted in an increase in the evenness (see Fig. 2, top right).

The post-disturbance evenness seems to drift towards a different state, which likely mirrors a different community structure. We also consider the correlation function, defined as the average over species of the autocorrelation function of each species’ abundance time series, i.e. 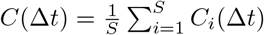 and 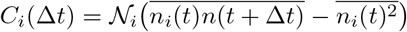, with 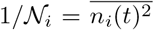, where an overline denotes time integration. We can estimate the correlation times of the system as 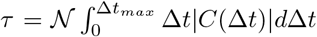, with 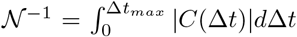 finding 8.88y for T0, 8.37y for T1, 8.55y for T2 and 8.82y for T3 (see Fig. 2, bottom panels). Temporal patterns are similar across different plot types, i.e. across a range of disturbance intensity. For this reason, we will focus on data collected from T3 plots.

## RESULTS AND DISCUSSION

We calculate the mean abundance from the empirical data and then use a two-fold Bayesian inference protocol [44] for the fit to (1): first, we fit simultaneously all the different typologies of perturbed plots (thus obtaining one set of parameters common to all plots). The parameters determined with this approach reflect the coarse behavior of the ecosystem following a perturbation. Secondly, we fit each plot individually while requiring that the population size returns asymptotically to pre-disturbance level. In this situation, plots are recognized as being impacted by separate disturbances. The comparison of successive iterations of the two approaches allows to guess sensible priors for parameters and latent initial conditions. The result of these approaches is a posterior distribution for all parameters and latent initial conditions. Notice that in the first case all mean abundances return to the same value, which is determined by the parameters of the fit. This may be different in general from the values of the pre-disturbance mean abundances. This is because we have assumed that each plot type has the same (effective) carrying capacity, which is, of course, only an approximation. In any case, the set of parameters obtained by fitting all plots together is compatible with those found by fitting the plots individually. Fig. 3 shows that the model is able to capture the empirical behavior quite well under different disturbance conditions.

The extension to multi-species models has two alternative formulations: a neutral model with noise and a high-dimensional model with heterogeneous interactions.

Each independent realization of the neutral model represents the dynamics of a single species. To obtain the dynamics of the full ecosystem, the initial conditions of the population size *n* are sampled from the empirical distribution of species’ abundances immediately after the perturbation. The model was simulated a number of times much larger than *S*. Conversely, the non-neutral model evolves all *S* species abundances simultaneously, encoding variability through heterogeneous species-species interactions. More details can be found in Appendix C.

To fit the models to data we followed two different approaches. Following the method of Approximate Bayesian Computation (ABC) [45, 46] the neutral model has been fitted to time-varying statistics extracted from data. Specifically, we computed numerically mean and variance of species abundances from the model and fit them to those found in data. The non-neutral model, in addition to the mean abundance data, has been calibrated from bootstrapped abundance data at stationarity. When it comes to fitting, the non-neutral model requires fewer empirical data compared to the neutral one, which instead relies on population time series for all species.

Once the multi-species models are calibrated from the time series, we can compute temporal community structure patterns that can be directly compared to the empirical patterns with no further fitting. We will consider the predictions for time correlation function, the temporal Taylor Law, the evenness and the evolution of the species abundance distribution (SAD).

As shown in Fig. 2, and from the correlation times computed, the empirical correlation functions of species abundances do not show a significant change as we consider different disturbance magnitudes. Therefore, it is likely that this pattern is driven by endogenous biological processes which are not sensitive to external perturbations. We find that both models predict a correlation that is compatible with the one observed in the data (see Fig. 4). The non-neutral model gives a better prediction of the magnitude, whilst the neutral model leads to a closer estimate of the correlation time (8.8*y* data, 7.9*y* neutral model). However, the empirical temporal correlation function is not sufficient to discriminate between neutral and non-neutral models.

**FIG. 4:**
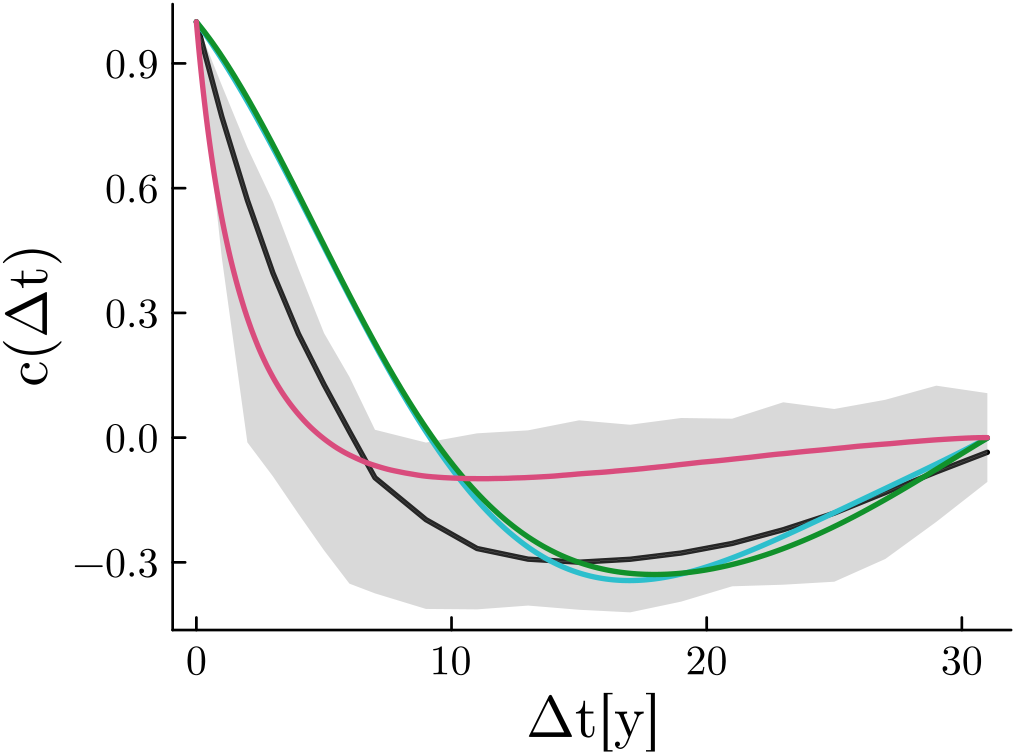
Temporal correlation function for the empirical data (black solid line with envelope given by 5 % and 95 % quantiles), non-neutral model (green: uncorrelated (*γ* = 0) couplings, cyan: anticorrelated (*γ* = ‐1) couplings) and neutral model (magenta).

Secondly, we have investigated the effects of the perturbations on the temporal Taylor Power Law (tTPL) [47, 48], which establishes a power law relation between the mean and the variance of abundances, namely

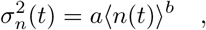

where 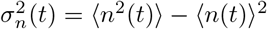 and *a, b* are two positive constants. We found that *b* is close to 2 (see Tab. V). This relation holds throughout post-disturbance dynamics. Given that the neutral model was calibrated directly on mean and variance of species abundances, it comes with no surprise that it can reproduce quite well the tTPL found in the data. On the other hand, the non-neutral model only used community information from bootstrapped time series and initial conditions. It is noteworthy that it can also replicate the tTPL, hinting at its ability to better describe community structure. Compared to the neutral model, we notice that the non-neutral deterministic dynamics with uncorrelated couplings 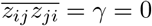 in (7)) accurately captures the power law exponent of the tTPL (see Fig. 5, left).

**FIG. 5:**
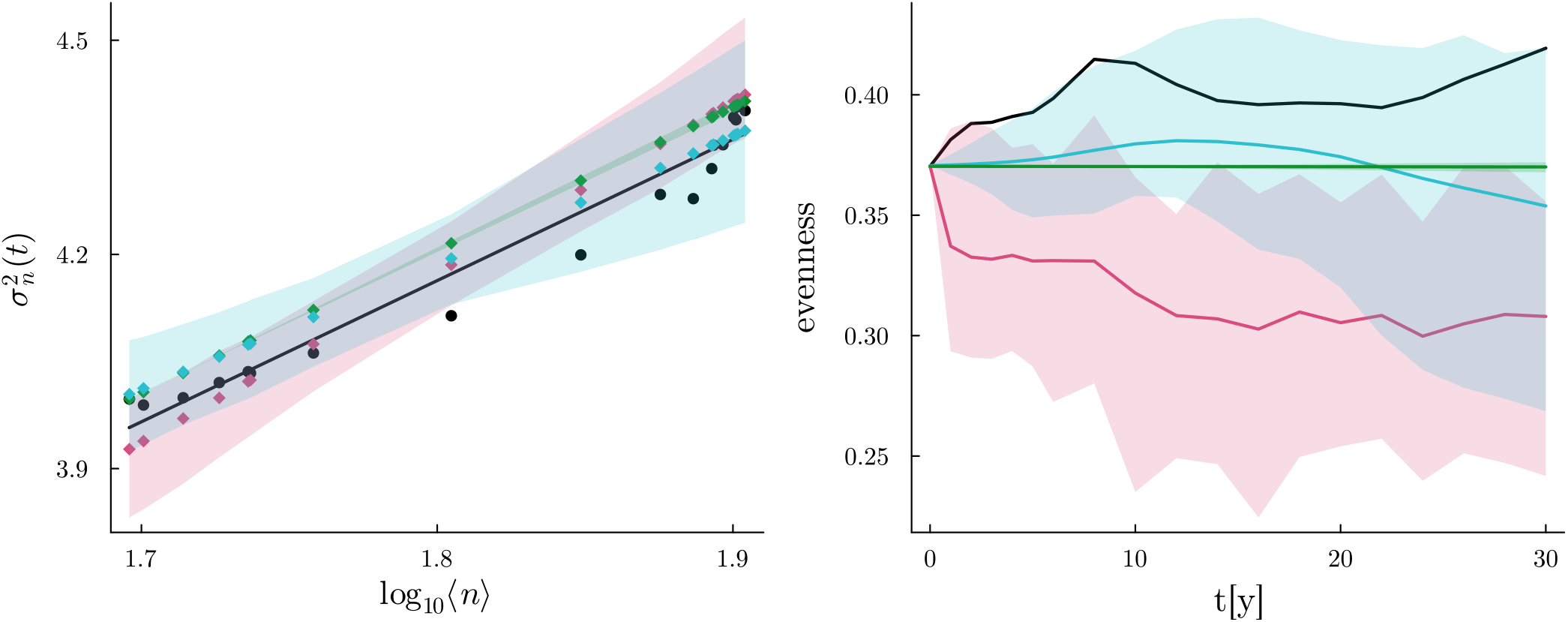
Temporal Taylor Power Law (left) and evenness (right) for empirical data (black), neutral model (magenta) and non-neutral model (green for uncorrelated couplings, cyan for anticorrelated couplings); envelopes are given by 5 % and 95 % quantiles obtained with 100 realizations of the models. Predicted values for the tTPL parameters are reported in Tab V.

In ecology, evenness [42] refers to the uniformity in the distribution of individuals among different species. High even-ness implies that species have comparable abundances, whereas low evenness suggests that a few species dominate the community. It is a crucial attribute of ecological communities, offering insights into the structure, function, and health of ecosystems, and facilitating their comparison. The definition we adopted here is independent of the total number of species, thus distinguishing community structure from species richness as separate aspects of biodiversity [49]. It has been shown that evenness can be a useful indicator for quantifying the detrimental influence of human-driven biodiversity depletion on primary productivity [43]. More broadly, evenness has been recognized to influence ecological functions, as well as temporal and spatial community variability, invadibility, and metacommunity dynamics [50].

The culling treatments carried out in the course of the PDEx smoothed out the differences in abundances between rare and commmon species, causing an increase in empirical post-disturbance evenness. Furthermore, over the following period, the community appears to remain in a stationary state which is different from the initial one, as signaled by the stable levels of evenness. It is interesting to notice that the level of evenness reached by the community after disturbance is largely independent of the magnitude of the disturbance. Fig. 5 shows that the non-neutral model predicts an evenness that is in good agreement with the observed data, thus indicating a superior ability in capturing community structure properties with respect to the neutral model. This occurs when the non-neutral model is endowed with interaction couplings which are anticorrelated 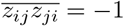. The physics literature on disordered ecological systems has shown that models of interacting species become less stable as the heterogeneity of interaction couplings increases, while their anticorrelation tends to make the system more stable [51, 52]. Consequently, if *γ* assumes negative values, the model remains stable even when heterogeneity is relatively strong. This result suggests that anticorrelated heterogeneity may play an important role in the dynamics of the system and better represents the community structure after a disturbance. On the contrary, as a results of species’ independence, the neutral model predicts much shorter relaxation times and smaller evenness compared to those observed. This suggests that the neutral framework, at least in the current formulation, is unable to capture important characteristics of the community. It is likely that during the recovery process the interactions between species play an important role, and the neutral assumption is simply too rough of an approximation to describe the dynamics after disturbance. This is in contrast to what happens at stationarity, where neutral models can explain quite well several patterns [19, 53–55].

## CONCLUSIONS

We outlined the key features of a consumer-resource model and connected it to data on transient population dynamics in a disturbed forest ecosystem. Initially, the model was fitted to average species abundances. To represent individual species explicitly, we extended the model into a multi-species framework using two approaches: a neutral model incorporating stochastic effects and a non-neutral one with heterogeneous couplings.

The non-neutral model’s predictions were observed to align more closely with the data than the predictions from the neutral one. This suggests that heterogeneity in species interactions helps to recover from a perturbation. This emerges more clearly from the behavior of the evenness. Also, because post-disturbance levels of evenness are relatively higher, this may indicate that the ecosystem has finally reached a long-lived structure different from the initial one, albeit the number of abundant species remained unaltered. The community may have used resources more complementarily, leading to increased plant recruitment [43].

It is interesting to note that the non-neutral model requires significantly less empirical information for calibration compared to the neutral one: estimating the total number of trees and species necessary for understanding mean abundance dynamics can be done more quickly than through individual tree censi, allowing for more cost-effective ecological monitoring and forecasting.

On a theoretical ground, instead, these results seem to caution against the naїve use of neutral models for ecosystems far from steady states. Indeed, models that are neutral and focus solely on demographic stochasticity often predict longer extinction times for common species, and temporal fluctuations tend to be underestimated [56, 57].

In future works, it will be useful to extend the model to explicitly represent space, thus looking at whether spatial data (which has not been used in this work) can lead to better insights and predictions. An extension of this study to transient dynamics during press perturbations is also of relevant interest, because it allows to assess the health of forest ecosystems in scenarios of global climate change. Conversely, to test the limits of applicability of our findings, it would be insightful to apply this framework to data coming from experiments in microbial ecology, because observations can be carried out in a more controlled manner and on time scales, relative to the intrinsic ones of the system, which are much longer than those achievable in forest ecosystems.

## Appendix A: (S,R)-Macarthur Model reduction to non-neutral model with R=1

We consider a system of S consumers’ species and R resources’ types

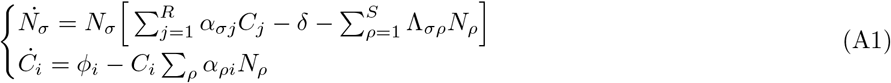

We now assume that the *total per capita* resource consumed is the same for all species, that is, defining *c* = (1*/R*) Σ _*j*_ *C*_*j*_, we assume ∀*σ* = 1, …, *S*

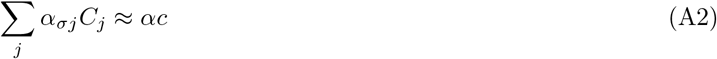

which is equivalent to assuming *α*_*σi*_ ≈ *α/R*. This allows us to rewrite (A1) in a coarse-grained form, namely

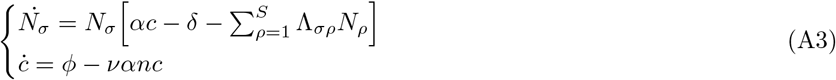

where 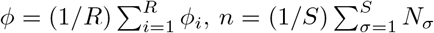 and *ν* = *S/R*. We fix the ratio *ν* to be equal to one for parsimony of parameters, thus recovering (6), which upon averaging over the *S* species of consumers is equivalent (on average) to (1).

## Appendix B: Approximations of the model of mean abundances

It is important to note that a quasi-stationary approximation, where *c* varies much faster than *n*, is not suitable for modeling the response to a pulse perturbation. By setting *ċ* = 0, we obtain an approximate solution to (1):

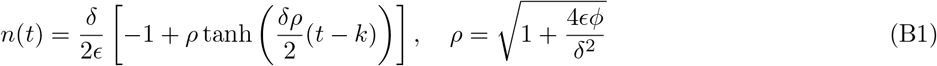

where *k* is uniquely determined by the initial condition *n*(0) = *n*_0_.

This solution increases monotonically and therefore cannot capture the overshooting of the data in Fig. 2. The variables *n* and *c* have to vary on the same temporal scale, which makes it difficult to introduce simple approximations. Alternatively, (1) can be linearised around its fixed point (2), finding the dynamics

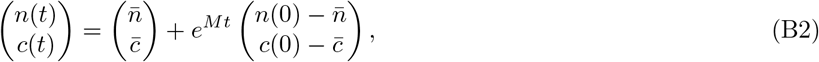

where *M* is the Jacobian matrix of (1) evaluated at the fixed point,

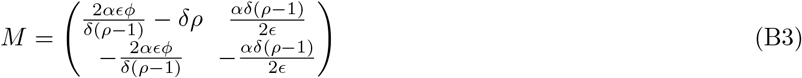

The exponential matrix in (B2) can be written explicitly; the solution of the linearized dynamics matches the full solution to a limited extent. We seek a better approximation to the full solution of (1) which we can still write down explicitly. We assume that the exact solution admits the Maclaurin expansions

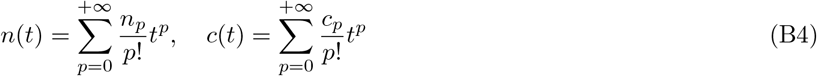

where 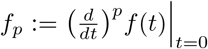 for *f* = *n, c*. Truncating the expansion (B4) at any given order will give a function which diverges as *t* → +∞.(To make progress, starting from the series for *n*(*t*) in (B4), we sum and subtract 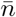 and multiply by 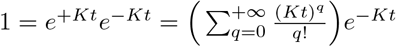 after collecting powers of *t*, we find

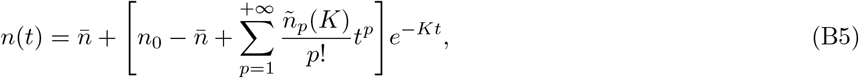

where we have defined the modified coefficients

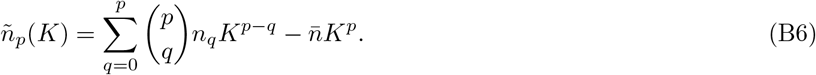

By suitably choosing *K >* 0 we can truncate this expansion at some order with a controlled error. Indeed if we neglect terms of order higher than L we introduce an error

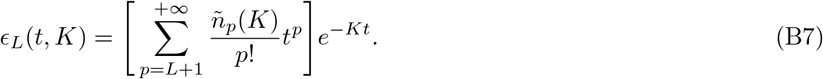

K should be taken so that the supremum of ∣*ϵ*_*L*_(*t, K*) ∣ as a function of time is minimized. In Fig. 6 we report a comparison between linearized dynamics and exponential cutoff approximation of the full solution.

**FIG. 6:**
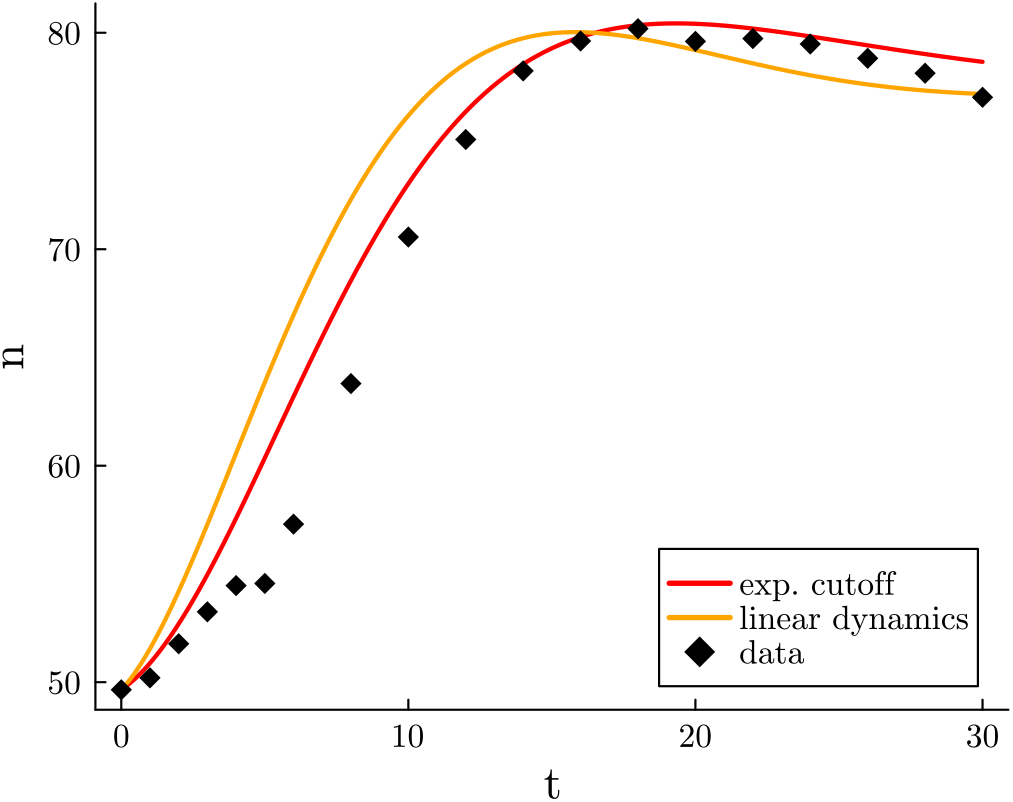
Fourth-order exponential cutoff approximation (red) and solution of linear dynamics (B2) (solid orange) compared to T3 data (black diamonds), with parameters found inferred as reported in Tab. II. For the cutoff approximation, *K* = 0.271 has been found through best fit; the closest natural time scale of the model is *ϵϕ/δ* = 0.309.

## Appendix C: Simulation and fit of neutral model

The system of stochastic equations (5) is interpreted with the Itô prescription of stochastic integrals [58]; reflecting boundary conditions are implemented through the transformation *x*→|*x*| applied to both *n* and *c* after each time step. The model has been integrated numerically according to the Euler-Maruyama scheme, with time step *dt* = 10^*‐*2^. We carried out a form of Approximate Bayesian Computation, in which parameters were selected to get the best fit of time series of the mean and variance of abundances obtained from the model to those extracted from data. During the fit, parameters were sampled from independent uniform distributions chosen empirically so that the model agreed with data; the best fit was thus obtained for parameters which minimized the square loss with respect to data. When running the model, the initial condition *n*_0_ has been sampled from the empirical distribution of abundances of *S* species in the year 1988, immediately after the disturbance has taken full effect in terms of death of individual trees. The sampling of initial conditions from the same distribution is the only link between different realizations of the model, which are otherwise completely independent from each other. At each step of the fit, we generated a number of single-species trajectories *N* ⪆ *S*· 10^2^ in order to obtain time series for mean and variance of abundances sufficiently smooth to be compared to data, where *S* is the number of species in the ecosystem.

Synthetic data used to derive pattern predictions have been generated by 100 runs of the model.

## Appendix D: Simulation and fit of mean abundance model and non-neutral model

To simulate ODE models we used Julia packages `DifferentialEquations.jl` for numerical integration, and `Turing.jl` for inference; at each step of the Monte Carlo Markov Chain algorithm used for inference, parameters have been sampled using the No-U-Turn Sampler (NUTS). For the mean abundance model (1), we chose independent uniform priors for each parameter. For the non-neutral model (6), all parameters other than *σ* have been fixed to the mean values of the posteriors obtained through inference on (1), and we chose a uniform prior distribution for *σ, σ* ∼ *U*_[*σ*_*‐,σ*+]. Since, for each value of *σ*, the solution of (6) is more likely to grow indefinitely as *γ* increases, we took *σ*^+^|_*γ*=*‐*1_ *> σ*^+^|_*γ*=0_. The posterior distribution for *σ* has been inferred through bootstrapping as follows. For species *i*, the mean and variance of abundance *µ*_*i*_ and *σ*_*i*_ before distrubance have been used to generate a time series of i.i.d random abundances with normal distribution, *n*_*t*_∼𝒩 (*µ*_*i*_, *σ*_*i*_). These time series have been used to generate a trajectory of evenness, to which evenness predicted by the model has been compared for different values of *σ* during inference.

Synthetic data used to derive pattern predictions have been generated by 100 runs of the model.

## Appendix E: Additional material: images, tables of fit parameters

**FIG. 7:**
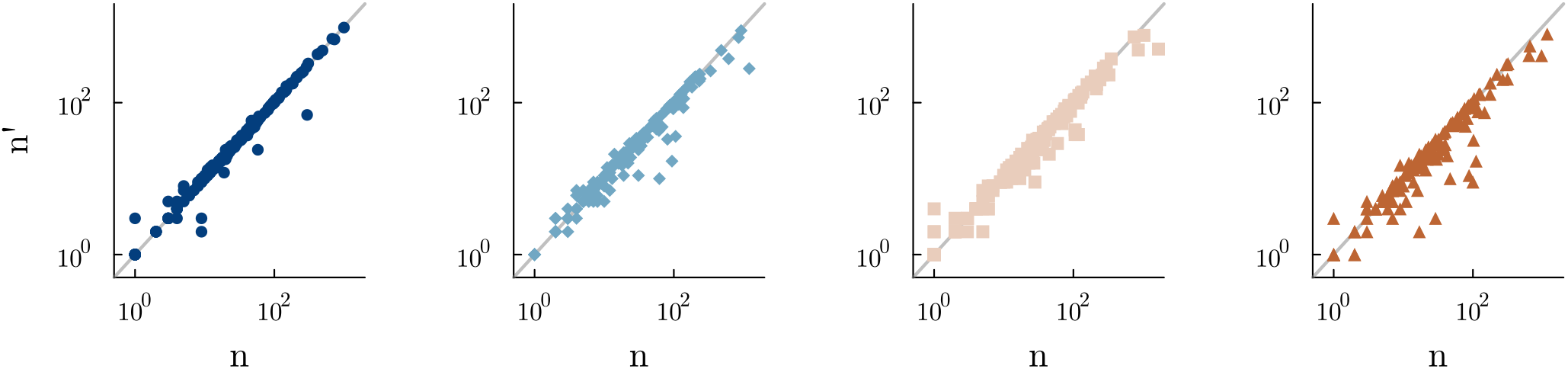
Distribution of perturbation across species: *n* and *n*^*′*^ denote respectively abundances of each species before and after treatment. Plot types are, from left to right, T0, T1, T2 and T3.

**TABLE II:** Parameters obtained from inference individually on different plot types (T1,T2,T3 columns) and simultaneously on all plots (Global column). When fitting individal plot types, convergence to steady state has been imposed by requiring a return to pre-perturbation mean abundances at time *t*_*st*_ = 500; resulting parameters are largely indepdendent on *t*_*st*_, provided *t*_*st*_ ≫ 30.

| Parameter | T1 | T2 | T3 | Global |
| --- | --- | --- | --- | --- |
| $\alpha$ | $(2.038 \pm 0.421) \times 10^{-3}$ | $(1.963 \pm 0.373) \times 10^{-3}$ | $(1.899 \pm 0.583) \times 10^{-3}$ | $(6.862 \pm 0.981) \times 10^{-4}$ |
| $\delta$ | $0.230 \pm 0.033$ | $0.197 \pm 0.028$ | $0.102 \pm 0.036$ | $0.072 \pm 0.023$ |
| $\epsilon$ | $(6.412 \pm 4.813) \times 10^{-4}$ | $(1.050 \pm 0.480) \times 10^{-3}$ | $(1.731 \pm 0.724) \times 10^{-3}$ | $(1.197 \pm 1.007) \times 10^{-4}$ |
| $\phi$ | $17.395 \pm 2.180$ | $19.471 \pm 2.602$ | $18.284 \pm 4.284$ | $4.381 \pm 1.456$ |

**TABLE III:** Parameters obtained from fitting the neutral model to data as described in the main text.

| Parameter | $\alpha$ | $\delta$ | $\epsilon$ | $\phi$ | $\sigma_n$ | $\sigma_c$ |
| --- | --- | --- | --- | --- | --- | --- |
| Value | $2.91 \times 10^{-4}$ | $1.20 \times 10^{-2}$ | $8.91 \times 10^{-4}$ | 24.2 | 9.67 | 0.412 |

**TABLE IV:** Interaction heterogeneity *σ* for the non-neutral model in the two cases of uncorrelated (*γ* = 0) and anticorrelated (*γ* = ‐1) interaction couplings.

| Parameter | $\sigma(\gamma = -1)$ | $\sigma(\gamma = 0)$ |
| --- | --- | --- |
| Value | $(6.87 \pm 3.95) \times 10^{-4}$ | $(7.01 \pm 39.2) \times 10^{-6}$ |

**TABLE V:** Predicted parameters of the tTPL in Fig. 5 obtained with a linear regression of the curve log 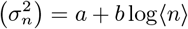.

| Model | Data | Neutral | Non-neutral ($\gamma = -1$) | Non-neutral ($\gamma = 0$) |
| --- | --- | --- | --- | --- |
| **a** | 1.388 | $-1.06 \pm 1.91$ | $3.35 \pm 1.81$ | $1.41 \pm 0.166$ |
| **b** | 1.978 | $2.56 \pm 0.468$ | $1.52 \pm 0.448$ | $1.997 \pm 0.042$ |

## References

[1] L. E. Frelich, “Forest dynamics and disturbance regimes: Studies from temperate evergreen-deciduous forests,” (2002).

[2] G. F. Peterken, “Structural dynamics of forest stands and natural processes,” in The Forests Handbook, Volume 1 (Black-well Science Ltd) p. 83–104.

[3] T. Mazor, C. Doropoulos, F. Schwarzmueller, D. W. Gladish, N. Kumaran, K. Merkel, M. Di Marco, and V. Gagic, Nature Ecology & Evolution 2, 1071–1074 (2018).

[4] E. A. Bender, T. J. Case, and M. E. Gilpin, Ecology 65, 1–13 (1984).

[5] A. Jentsch and P. White, Ecology 100 (2019), 10.1002/ecy.2734.

[6] J. Maximillian, M. Brusseau, E. Glenn, and A. Matthias, “Pollution and environmental perturbations in the global system,” in Environmental and Pollution Science (Elsevier, 2019) p. 457–476.

[7] I. Donohue, O. L. Petchey, J. M. Montoya, A. L. Jackson, L. McNally, M. Viana, K. Healy, M. Lurgi, N. E. O’Connor, and M. C. Emmerson, Ecology Letters 16, 421–429 (2013).

[8] J.-F. Arnoldi, M. Loreau, and B. Haegeman, Journal of Theoretical Biology 389, 47–59 (2016).

[9] V. Radchuk, F. D. Laender, J. S. Cabral, I. Boulangeat, M. Crawford, F. Bohn, J. D. Raedt, C. Scherer, J. Svenning, K. Thonicke, F. M. Schurr, V. Grimm, and S. Kramer-Schadt, Ecology Letters 22, 674–684 (2019).

[10] S. Kéfi, V. Domínguez-García, I. Donohue, C. Fontaine, E. Thébault, and V. Dakos, Ecology Letters 22, 1349–1356 (2019).

[11] A. T. Clark, J. Arnoldi, Y. R. Zelnik, G. Barabas, D. Hodapp, C. Karakoç, S. König, V. Radchuk, I. Donohue, A. Huth, C. Jacquet, C. de Mazancourt, A. Mentges, D. Nothaaß, L. G. Shoemaker, F. Taubert, T. Wiegand, S. Wang, J. M. Chase, M. Loreau, and S. Harpole, Ecology Letters 24, 1474–1486 (2021).

[12] I. Stott, Methods in Ecology and Evolution 7, 666–678 (2016).

[13] D. N. Koons, D. T. Iles, and I. Stott, “Transient analyses of population dynamics using matrix projection models,” in Demographic Methods across the Tree of Life (Oxford University PressOxford, 2021) p. 197–212.

[14] “Guyafor, database of the french guiana permanent plot network, cirad-cnrs-onf,” http://www.ecofog.gf/spip.php?article364.

[15] “Paracou disturbance experiment webpage,” https://paracou.cirad.fr/website/experimental-design/disturbance-experiment.

[16] C. Baraloto, B. Hérault, C. E. T. Paine, H. Massot, L. Blanc, D. Bonal, J. Molino, E. A. Nicolini, and D. Sabatier, Journal of Applied Ecology 49, 861–870 (2012).

[17] P. Sist, L. Blanc, L. Mazzei, C. Baraloto, and R. Aussenac, Bois et Forêts des Tropiques (4), 41 (2012).

[18] S. P. Hubbell, “The unified neutral theory of biodiversity and biogeography,” (2001).

[19] S. Azaele, S. Suweis, J. Grilli, I. Volkov, J. R. Banavar, and A. Maritan, Reviews of Modern Physics 88, 035003 (2016).

[20] I. C. Barrett, A. R. McIntosh, and H. J. Warburton, Community Ecology 24, 257–269 (2023).

[21] S. R. Supp and S. K. M. Ernest, Ecology 95, 1717–1723 (2014).

[22] B. Worm, H. K. Lotze, H. Hillebrand, and U. Sommer, Nature 417, 848–851 (2002).

[23] J. Wootton, The American Naturalist 152, 803–825 (1998).

[24] Y. Yang, K. Z. Coyte, K. R. Foster, and A. Li, Nature Communications 14 (2023), 10.1038/s41467-023-42580-0.

[25] T. Galla, EPL (Europhysics Letters) 123, 48004 (2018).

[26] A. R. Batista-Tomás, A. De Martino, and R. Mulet, Chaos: An Interdisciplinary Journal of Nonlinear Science 31 (2021), 10.1063/5.0046972.

[27] D. Gupta, S. Garlaschi, S. Suweis, S. Azaele, and A. Maritan, Physical Review Letters 127 (2021), 10.1103/phys-revlett.127.208101.

[28] T. Arnoulx de Pirey and G. Bunin, Physical Review X 14 (2024), 10.1103/physrevx.14.011037.

[29] E. Blumenthal, J. W. Rocks, and P. Mehta, Physical Review Letters 132 (2024), 10.1103/physrevlett.132.127401.

[30] S. P. Hubbell, Functional Ecology 19, 166–172 (2005).

[31] S. Suweis, F. Ferraro, C. Grilletta, S. Azaele, and A. Maritan, “Generalized lotka-volterra systems with time correlated stochastic interactions,” (2024), 2307.02851 [q-bio.PE].

[32] R. Lande, Oikos 83, 353 (1998).

[33] R. Lande, S. Engen, and B.-E. Saether, “Stochastic population dynamics in ecology and conservation,” London, England (2003).

[34] S. Azaele, S. Pigolotti, J. R. Banavar, and A. Maritan, Nature 444, 926–928 (2006).

[35] A. Traulsen, J. C. Claussen, and C. Hauert, Physical Review E 85 (2012), 10.1103/physreve.85.041901.

[36] D. A. Vasseur and P. Yodzis, Ecology 85, 1146–1152 (2004).

[37] P. Chesson, Theoretical Population Biology 37, 26–38 (1990).

[38] P. Chesson, Annual Review of Ecology and Systematics 31, 343–366 (2000).

[39] C. Daniel, E. Allan, H. Saiz, and O. Godoy, Ecology Letters 27 (2024), 10.1111/ele.14425.

[40] S. Gourlet-Fleury, J.-M. J.-M. Guehl, and O. Laroussinie, “Ecology and management of a neotropical rainforest. Lessons drawn from Paracou, a long-term experimental research site in French Guiana,” (2004), 350 p. pp.

[41] J. Gauthier and N. Derome, mSphere 6 (2021), 10.1128/msphere.01019-20.

[42] B. Smith and J. B. Wilson, Oikos 76, 70 (1996).

[43] B. J. Wilsey and C. Potvin, Ecology 81, 887–892 (2000).

[44] A. Gelman, J. B. Carlin, H. S. Stern, D. B. Dunson, A. Vehtari, and D. B. Rubin, “Bayesian data analysis,” Philadelphia, PA (2013).

[45] P. J. Diggle and R. J. Gratton, Journal of the Royal Statistical Society Series B: Statistical Methodology 46, 193–212 (1984).

[46] M. Sunnåker, A. G. Busetto, E. Numminen, J. Corander, M. Foll, and C. Dessimoz, PLoS Computational Biology 9, e1002803 (2013).

[47] M. R. D. Cobain, M. Brede, and C. N. Trueman, Journal of Animal Ecology 88, 290–301 (2018).

[48] H. Kojima, Y. Mitsui, and T. Ikegami, Chaos: An Interdisciplinary Journal of Nonlinear Science 31 (2021), 10.1063/5.0036892.

[49] L. Jost, Diversity 2, 207–232 (2010).

[50] H. Hillebrand, D. M. Bennett, and M. W. Cadotte, Ecology 89, 1510–1520 (2008).

[51] S. Allesina and S. Tang, Nature 483, 205–208 (2012).

[52] G. Bunin, Phys. Rev. E 95, 042414 (2017).

[53] I. Volkov, J. R. Banavar, S. P. Hubbell, and A. Maritan, Nature 424, 1035–1037 (2003).

[54] I. Volkov, J. R. Banavar, S. P. Hubbell, and A. Maritan, Nature 450, 45–49 (2007).

[55] F. Peruzzo, M. Mobilia, and S. Azaele, Physical Review X 10, 011032 (2020).

[56] C. M. Mutshinda, R. B. O’Hara, and I. P. Woiwod, Functional Ecology 22, 340–347 (2007).

[57] R. E. Ricklefs, Ecology 87, 1424–1431 (2006).

[58] C. W. Gardiner, “Handbook of stochastic methods for physics, chemistry and the natural sciences,” Berlin (2004), xviii–415 pp.

